# Plant architecture as a tool to mitigate late blight: significant but variable contributions of erect, aerated potato canopies over six years of field trials

**DOI:** 10.64898/2026.09.22.753506

**Authors:** Claudine Pasco, Melen Leclerc, Christophe Langrume, Bruno Marquer, Didier Andrivon

## Abstract

To reduce reliance on pesticides in potato production, alternative partial control methods are needed, yet their efficacy and reliability are seldom quantified over multiple years. Canopy architecture is a potential lever against late blight, caused by *Phytophthora infestans*, a pathogen whose infection and development are highly driven by humidity and temperature. However, its practical value under field conditions in pure crop stands remains poorly documented. We compared two commercial potato cultivars with contrasting canopy architectures but similar leaf tissue susceptibility to *P. infestans*, in replicated field trials conducted over six years (2011–2016) under irrigation, and over three of these years also without irrigation. Disease and canopy development (height, closure, leaf area, stem number) were monitored throughout the growing seasons and analysed using non-linear growth models. Monalisa, an erect and ramose cultivar, consistently slowed epidemic progress relative to Bintje, a semi-erect and leafy cultivar. This effect was significant in every non-irrigated trial but in only half of the irrigated ones. No single canopy trait explained this suppression across all years and conditions. However, canopy closure at the time of inoculation was the most consistent correlate, supporting the hypothesis of a microclimate-mediated effect. Disease reduction seldom translated into a yield benefit, except under irrigation, where Monalisa occasionally outyielded Bintje. Our results confirm that canopy architecture can complement other levers within integrated late blight management. Nevertheless, its significant year-to-year variability and dependence on environmental conditions must be accounted for if it is to be deployed as a reliable control method.

## 1 Introduction

Reducing the use of synthetic pesticides for crop protection now stands high on the political agenda of many countries, notably within the European Union (Jacquet et al., 2024). Reaching this goal however necessitates a major paradigm shift in the design and planning of plant production systems by incorporating plant protection among the strategic decisions rather than resorting to only tactical measures (when and what to spray?) (Lamichhane et al., 2016). This paradigm shift, in turn, requires the development and assessment of alternatives that mitigate disease in host crops, which can then be incorporated and integrated into redesigned cropping systems (Deguine et al., 2021; Jacquet et al., 2024). However, contrary to pesticides, most of these alternatives such as biocontrol agents, cultural practices, or host resistance tend to provide only partial and context dependent control (Andrivon et al., 2003; Bruce et al., 2017). Although the integration of the uncertainties associated to such partial control in epidemiological models would be useful for farmers and agronomists to better handle the risk of disease development and optimise the design of integrated plant production systems, their quantification requires considerable field data that rarely exist.

Plant-based levers for disease control, in particular qualitative and quantitative host resistance, remain the most popular alternative to pesticide use worldwide (Lannou, 2012). Although breeding for resistance has achieved considerable success in many crops, the rapid evolution of pathogens can undermine resistance durability and efficacy (Consortium, 2016). Further to resistance, plant or canopy architecture has long been recognised to influence pathogen development in host crops and can be used to mitigate epidemic spread (Calonnec et al., 2013; Tivoli et al., 2013). Plant and crop architecture, i.e. the density and spatial distribution of host organs within the canopy, is indeed known to impact microclimatic conditions (Richard et al., 2013; Vidal et al., 2017), in particular variables that are crucial for infection and dispersal of pathogens, and to act on the physiology and consequently on the susceptibility of host tissues (Robert et al., 2018; Motisi et al., 2019).

Plant or canopy architecture can be manipulated to reduce pathogen development in two main ways. First, some architectural traits are heritable and can be used in plant breeding programmes (Andrivon et al., 2013). This is particularly interesting when sources of genetic resistance are very limited (Ando et al., 2007). Second, plant and canopy architecture can be manipulated by human interventions before and during the cropping season (Costes et al., 2013; Mammeri et al., 2014). One can choose the spatial arrangements of plants at sowing or planting, choose the sowing date, use irrigation and fertilisation to impact plant growth or remove or rearrange spatially part of the plants (Levionnois et al., 2023).

In this study, we experimentally assess to what extent plant canopy can slow down the spread of potato late blight in field conditions. Caused by the oomycete *Phytophthora infestans*, late blight is the most damaging disease affecting potato (*Solanum tuberosum* L.) production globally (Haverkort et al., 2008; Acuña et al., 2023). Following its historical role in the Irish potato famine, the pathogen continues to cause substantial economic losses due to its rapid reproductive cycle, prolific sporulation, and adaptability to diverse environments. The use of fungicides and genetic resistance remain the cornerstone of disease development worldwide. However, the explosive demography of *P. infestans* still challenge its durable control (Cooke et al., 2011; Leesutthiphonchai et al., 2018). This underscores the need for diversified and integrated control strategies that reduce the reliance on a single method. While much attention has been paid to both quantitative and qualitative genetic resistances to late blight in potato, the potential for plant architecture to slow disease development in the field remains less explored. In theory, acting through plant architecture is particularly relevant for pathogens like *P. infestans*, which depend heavily on leaf wetness and humidity for infection and dispersal (Olanya et al., 2007; Launay et al., 2014). An architectural ideotype to limit the progress of potato late blight has already been proposed by Andrivon et al. (2013). The ideal potato would have long internode length, small leaflets and a low leaf area index (LAI), a low stem density and leaf area density (LAD), and slow canopy closure (Casadebaig et al., 2012). In the past, these traits have been strongly selected against in breeding programmes, as they tend to reduce potential yield. However, as the effects of architecture on potato late blight remain poorly documented in the literature, this ideotype was designed on postulated effects that still need to be confirmed by experimental data.

To isolate the contribution of architectural traits to epidemic development, we focus here on two commercial cultivars which exhibit similar levels of tissue susceptibility to *P. infestans* but different levels of field resistance, attributed to plant architectures more or less favourable to infection and epidemic spread. Over a six-year period, we conducted replicated field experiments under both irrigated and non-irrigated conditions to assess the consistency, magnitude, and agronomic relevance of architecture-driven disease suppression. Canopy and disease data acquired on the two cultivars were compared to evaluate the effect of architecture on the progression of late blight epidemics, and identify which architectural traits are associated with reduced disease development. By integrating original long-term field data, this study also enabled us to assess the variability of canopy traits and control efficacy through architecture. We finish by discussing how our results provides new insights into the epidemiological role of canopy structure, its potential as a complementary lever in integrated disease management strategies.

## 2 Materials and Methods

### 2.1 Plant material

Two commercial potato cultivars, Bintje and Monalisa, were selected for this study due to their contrasting canopy architectures and field resistance to late blight, but similar tissue susceptibility to *P. infestans*. Bintje is described in the French Catalogue of Potato Varieties (FN3PT – GNIS ARVALIS - Institut du végétal, 2018) as a moderately tall, semi-erect and rather leafy cultivar, susceptible to foliage blight (scored 3 on a 1-9 scale of increasing resistance), while Monalisa is described as a moderately tall, erect and ramose type, with a rather low susceptibility to foliage blight (scored 6 on the 1-9 scale of increasing resistance). Bintje canopies are therefore more closed and less aerated than those of Monalisa. Both cultivars belong to the same, semi-early maturity group.

### 2.2 Leaf tissue susceptibility test

We used a standard biotest protocol developed in our laboratory (Clément et al., 2010) to confirm the similar tissue susceptibility of the two cultivars to the late blight pathogen *P. infestans*. Forty detached leaflets of each cultivar were inoculated individually by depositing a 20*µ*L droplet containing about 1000 *P. infestans* sporangia (isolate 13P19.03, clonal lineage EU-06-A1) at the centre of each leaflet, placed abaxial face up. Inoculated leaflets were incubated in clear plastic boxes stored at 18 °C and 15 °C (night) with 16 h of daylight. Six days later, the lesion size was assessed on each leaflet by measuring minor and major radii of the lesion with a sliding caliper, and lesion area was calculated assuming an elliptical shape, which provides a good fit to actual size (Leclerc et al., 2019). Thereafter, sporangia were collected by washing each leaflet in 10 mL Isoton (Saline buffer; Beckman Coulter, Villepinte, France) and counted with a Coulter Z2 counter (Beckman Coulter with lower and upper thresholds of 10*µ*m and 20*µ*m respectively). Sporulation capacity (in sporangia per cm^2^ of lesion) was then calculated, and both lesion area and sporulation capacity data were statistically compared between cultivars using one-way analysis of variance (ANOVA).

### 2.3 Field trials

For the two cultivars, plant cover and disease development were assessed under field conditions over six consecutive years (2011–2016). Field trials were conducted at the INRAE experimental station La Motte in Le Rheu, France (48°06’N, 1°48’W). At this site the soil is overall silty with mean pH value of 6.69. Each experimental plot consisted of 10 rows of 15 certified seed tubers (size 35/45 mm), spaced 30 cm within rows and 70 cm between rows. Each year, both cultivars were grown in both unprotected or fungicide protected plots, the latter serving as controls. Fungicide protection followed standard local recommendations, though specific product details were not the primary focus of this study. In order to promote disease development, overhead sprinkler irrigation was applied during the six years. In addition, during three years (2012–2014), non-irrigated potato plots were also included in the trial design to compare highly favourable (irrigated) and less favourable conditions on canopy and disease development (Table S1).

As in this area, potato production is not common and inoculum pressure may be low, artificial inoculations were used to induce late blight epidemics. Inoculum was prepared by infecting Bintje leaflets with a virulent isolate of *P. infestans* from our laboratory collection. Sporulating detached leaflets, produced as described above for the susceptibility tests, were affixed to outer-row plants using clips and enclosed in transparent plastic bags for six days to ensure high humidity and facilitate pathogen establishment. Inoculations were carried out two months after planting (16–22 June) (Table S1).

In each plot, the canopy structure was characterised by measuring several traits on three randomly placed quadrats of one m^2^ each (Table S2). Canopy coverage, height and total leaf surface per plant were monitored, once or twice a week, between emergence and senescence, while the number of stems per plant and the yield were assessed after inoculation and harvest, respectively. Canopy coverage was estimated by calculating the proportion of green foliage in an azimuth image of the quadrat acquired with a standard digital camera. Canopy height was measured on 4 plants within each quadrat with a standard measuring ruler. Leaf surface per plant was obtained through a destructive sampling in protected plots. At each sampling date, two plants were collected and the total leaf surface was assessed with a planimeter (LI-3100C Leaf Area Meter, LI-COR) (Jumel et al., 2025). The number of stems per plant was counted in protected plots for 4 (2011–2013), 6 (2014–2015) or 9 (2016) plants at each sampling date. Total tuber yield per plot was determined by weighing the harvested daughter tubers of each experimental plot. Finally, late blight development was monitored throughout the epidemics by estimating the percentage of diseased foliage on each quadrat using a visual scoring scale (Pellé et al., 2005).

### 2.4 Statistical analysis and modelling

We considered two parsimonious non-linear models to analyse data over time. Canopy growth dynamics, characterised using canopy coverage, height and leaf surface per plant, was described using a double logistic function that captures expansion and senescence phases:

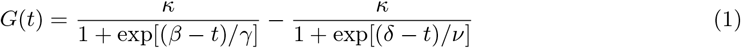

where *κ* is the upper asymptote, *β* and *δ* are respectively the inflection points for growth and senescence, *γ* and *v* are respectively inverse growth and senescence rates.

Disease progress curves were analysed using a logistic growth function:

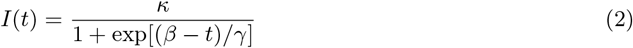

where the parameters have the same meaning of above.

To account for temperature effects on pathogen development and plant growth, all time-dependent processes were analysed relative to degree-days (base 0 °C) from plant emergence, assumed to occur 125 degree-days post-planting (MacKerron and Waister, 1985). The models were fitted to the data using non-linear least squares.

For disease development, we used the fitted curves to compute the area under the disease progress curve (AUDPC) between 0 and 900 degree-days using a trapezoidal integration (Jeger and Viljanen-Rollinson, 2001):

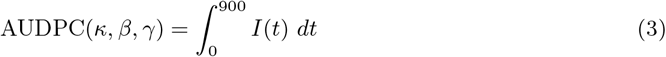

In addition, to compare the two cultivars we calculated the disease suppression benefit of Monalisa relative to Bintje as follows:

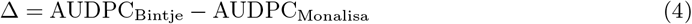

The effects of cultivar, year, irrigation, and their interactions on canopy traits, stem number, and yield were assessed using linear models and ANOVA. Differences in architectural traits and yield between cultivars were compared for each year and irrigation regime. Correlations between canopy traits at inoculation time and subsequent disease development were computed to evaluate potential explanatory variables for the observed differences in disease progress. All statistical analyses were performed in R (R Core Team, 2025).

## 3 Results

### 3.1 Leaf tissue susceptibility

Controlled-condition biotests on detached leaflets confirmed that both cultivars exhibited similar levels of susceptibility to *P. infestans*. No significant differences were observed in lesion size (*p* = 0.58), nor in sporulation capacity (*p* = 0.37) between Bintje and Monalisa (Fig. 1).

**Figure 1.**
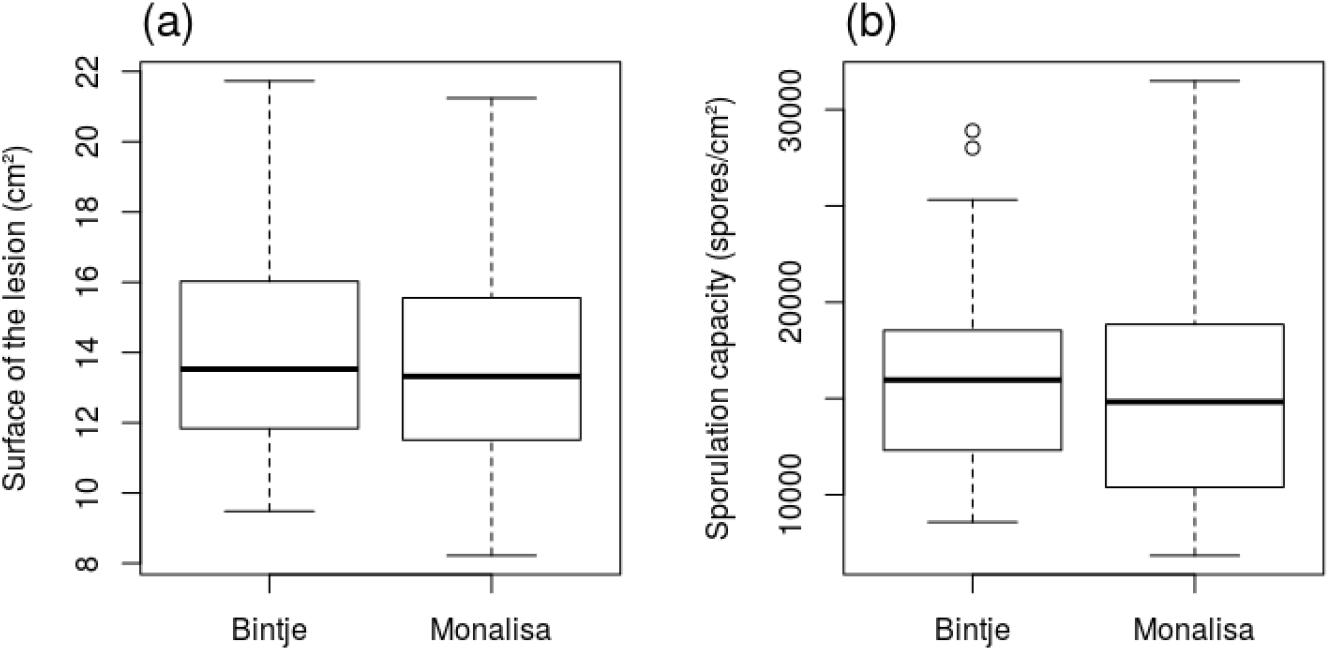
Comparison of the susceptibility of cultivars. The figures shows the distribution of a) the surface of lesions and b) the sporulation capacity, both obtained 6 days after inoculation.

### 3.2 Variation in canopy architecture

Canopy traits displayed substantial variation across years and between cultivars (Figs. 2-5). As initially expected, Monalisa typically developed taller plants with sparser foliage and fewer stems. On the contrary, Bintje showed more compact, dense and horizontally extensive growth. However, over the six years of trials, there are always years that contradict these general trends. For instance, Bintje had a higher leaf surface per plant than Monalisa except for the year 2014 where Monalisa had substantially dense foliage. Also, although Monalisa was usually taller than Bintje, the later was taller in 2013 and 2014.

**Figure 2.**
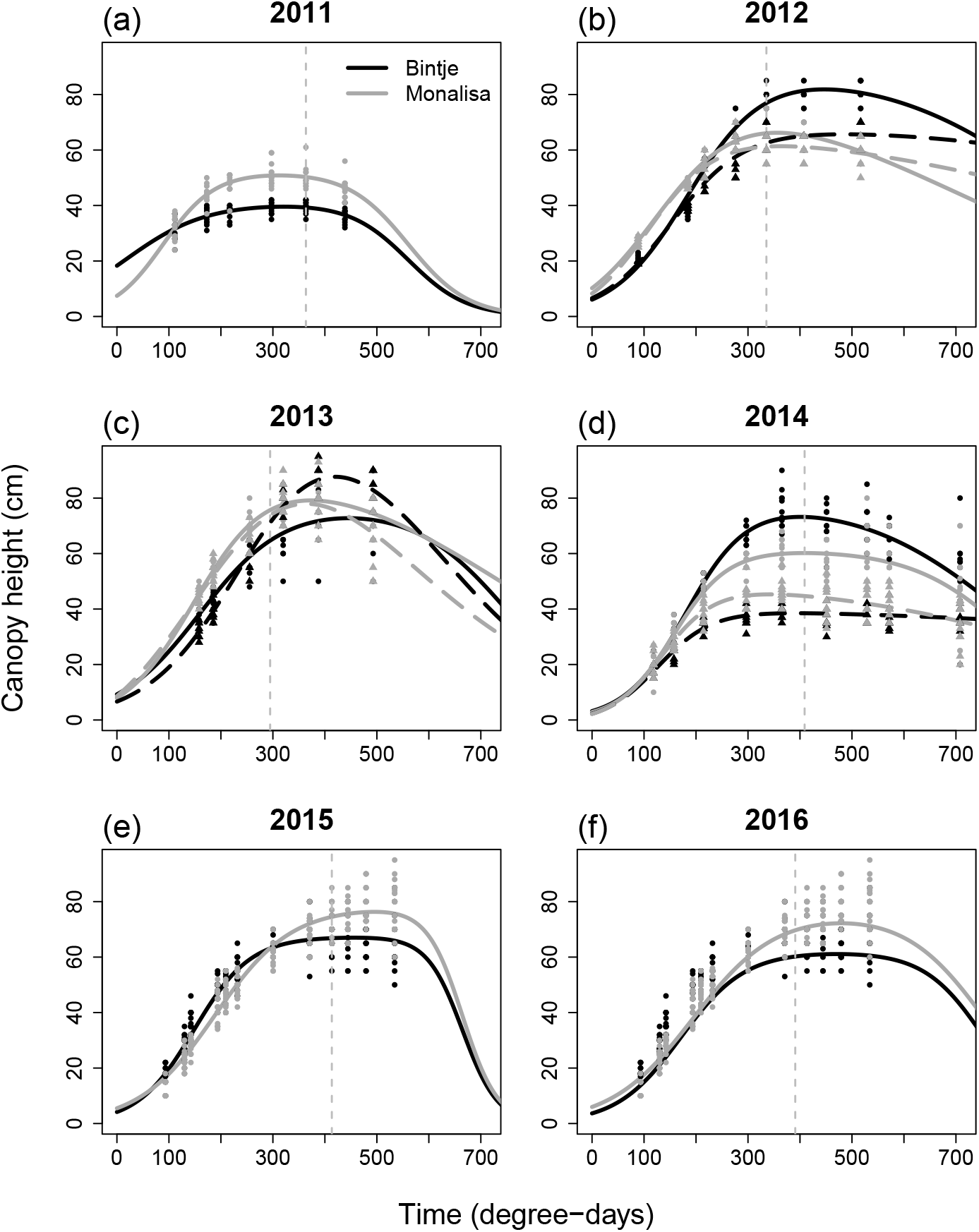
Temporal dynamics of canopy height. Observed data and estimated growth curves across six years of experiments. Black lines and dots represent the Bintje variety, while grey lines and dots represent Monalisa. Solid lines indicate disease progress in irrigated plots (2011–2016), and dashed lines (2012–2014) show disease development in non-irrigated plots. For each year, the inoculation time is marked by a vertical grey dotted line.

The double logistic model (eq. 1) allowed us to analyse the longitudinal data and capture the essential pattern of canopy height (Fig. 2 & Table S3), coverage (Fig. 3 & Table S4) and leaf surface per plant (Fig. 4 & Table S5). The ANOVA performed on stem numbers (Fig. 5) pointed out significant effects of cultivar, year, and their interaction underscoring temporal variability in morphological expression (p-values*<* 0.05) (Table S6). A post-hoc analysis through Tuckey’s tests revealed that the number of stems was significantly higher in Bintje only for years 2014, 2015 and 2016 (Table S7). The effect of irrigation on stem number was negligible, suggesting that this architectural trait is robust to water availability (p= 0.49).

**Figure 3.**
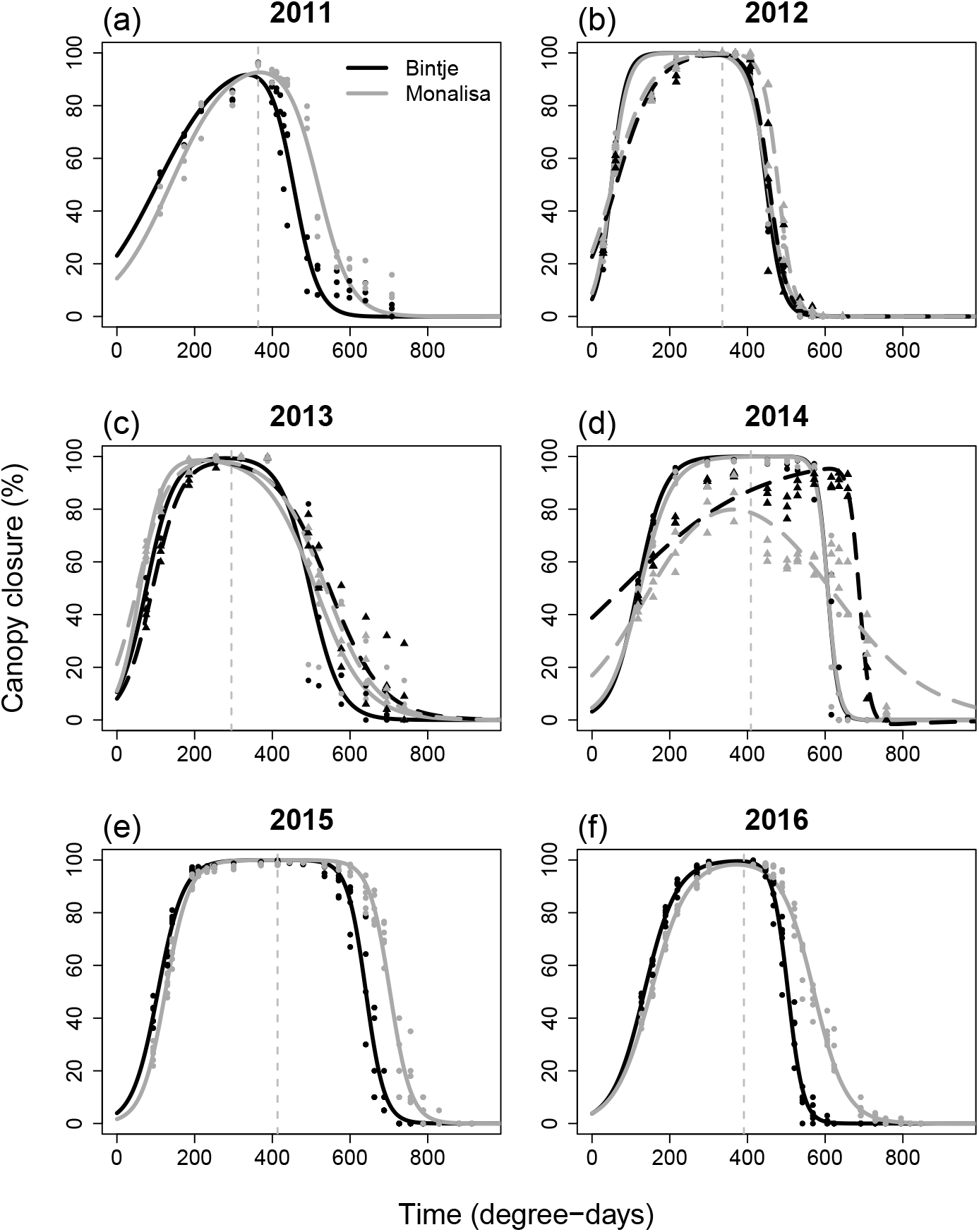
Temporal dynamics of canopy closure. Observed data and estimated growth curves across six years of experiments. Black lines and dots represent the Bintje variety, while grey lines and dots represent Monalisa. Solid lines indicate disease progress in irrigated plots (2011–2016), and dashed lines (2012–2014) show disease development in non-irrigated plots. For each year, the inoculation time is marked by a vertical grey dotted line.

**Figure 4.**
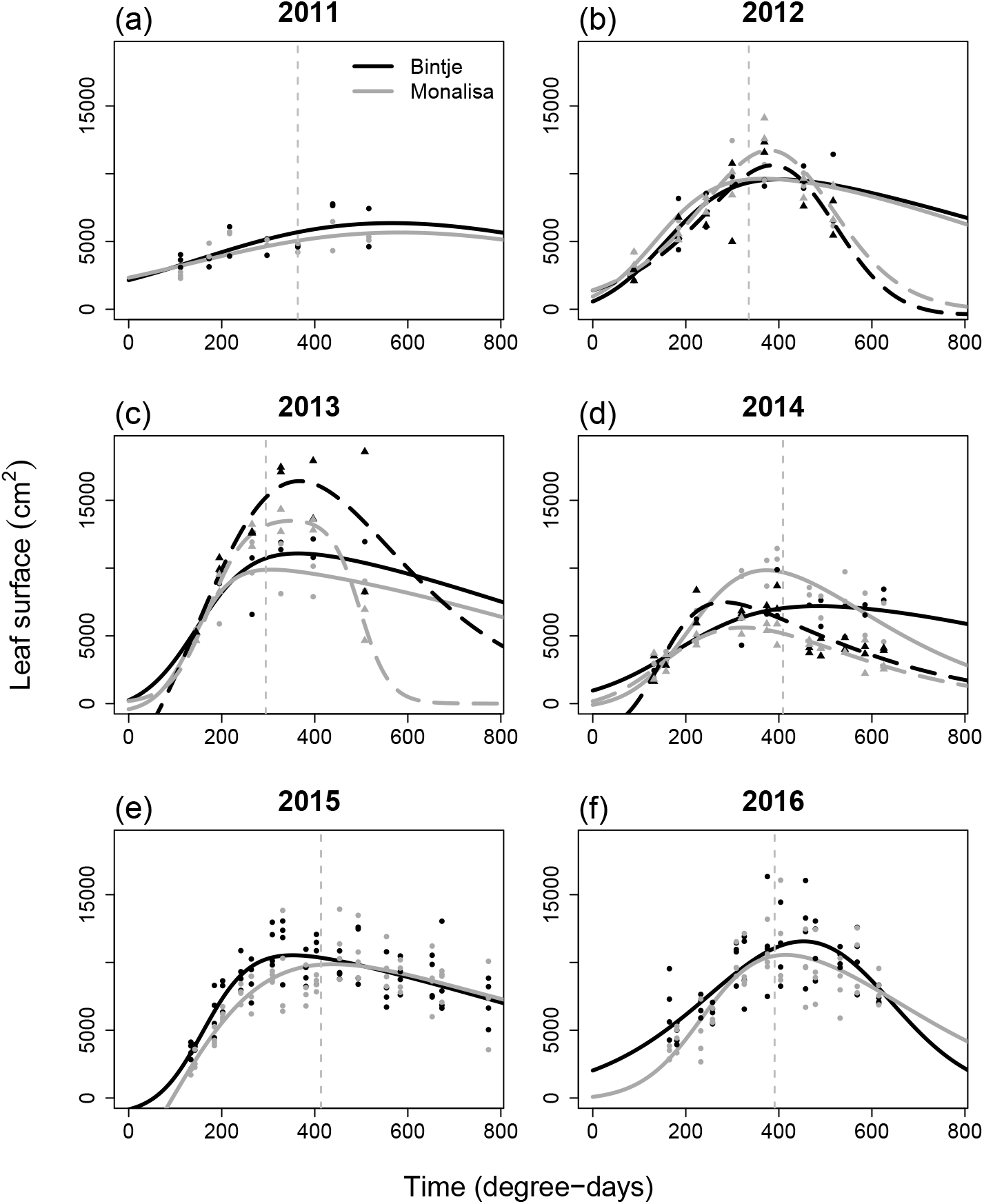
Temporal dynamics of leaf surface. Observed data and estimated growth curves across six years of experiments. Black lines and dots represent the Bintje variety, while grey lines and dots represent Monalisa. Solid lines indicate disease progress in irrigated plots (2011–2016), and dashed lines (2012–2014) show disease development in non-irrigated plots. For each year, the inoculation time is marked by a vertical grey dotted line.

**Figure 5.**
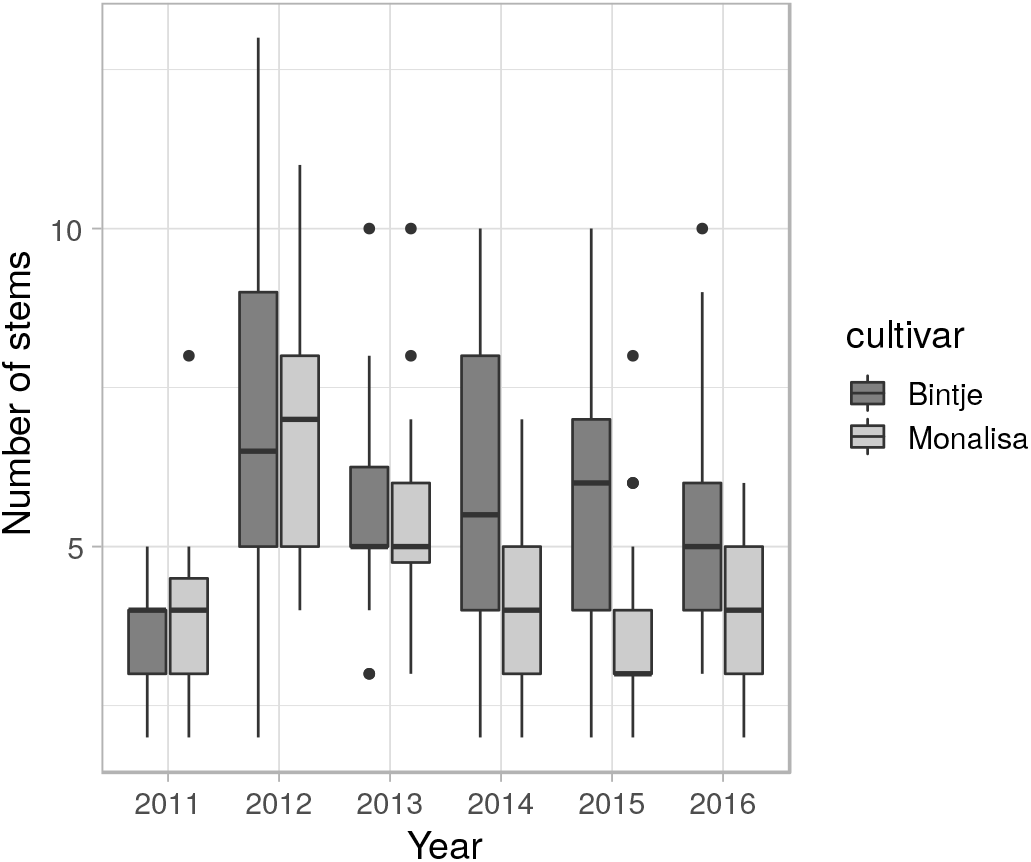
Number of stems of canopies across the six years. The Bintje variety is represented in dark grey, while Monalisa is in light grey.

The magnitude and timing of canopy closure, height development, and leaf surface area fluctuated strongly from year to year, and this variability was often greater than the inter-varietal one.

### 3.3 Disease progress in field trials

The logistic model (eq. 2) essentially captured the disease progress data (Fig. 6) and the fitting procedure allowed to estimate model parameters from the data (Table 1). Across the six-year experiments, disease progress curves revealed substantial year-to-year variability. However, two general trends emerged.

**Table 1.** Estimated parameters of the logistic model fitted to disease progress data. AUDPC was calculated by numerically integrating the estimated logistic curve between 0 and 900 degree-days. The p-value corresponds to a Fisher test that compared common and separated curves. The benefit Δ is the difference between AUDPC of Bintje and Monalisa.

| | Year | Cultivar | $\kappa$ | $\beta$ | $\gamma$ | AUDPC | p-value | Benefit $\Delta$ |
| --- | --- | --- | --- | --- | --- | --- | --- | --- |
| Irrigated |  |  |  |  |  |  |  |  |
|  | 2011 | Bintje | 95.8 | 459.8 | 25.1 | 42189.9 |  |  |
| | | Monalisa | 95.4 | 522.6 | 32.8 | 36022.3 | $< 10^{-15}$ | 6167.6 |
|  | 2012 | Bintje | 100.0 | 440.0 | 11.1 | 45985.8 |  |  |
|  |  | Monalisa | 98.7 | 444.5 | 8.3 | 44944.3 | 0.30 | 1041.5 |
|  | 2013 | Bintje | 100.0 | 480.1 | 7.9 | 41990.0 |  |  |
|  |  | Monalisa | 95.7 | 474.3 | 16.4 | 40743.7 | **0.07** | 1246.3 |
|  | 2014 | Bintje | 100.0 | 600.5 | 15.7 | 29950.0 |  |  |
|  |  | Monalisa | 98.9 | 599.2 | 11.3 | 29740.9 | 0.47 | 209.1 |
|  | 2015 | Bintje | 100.0 | 602.4 | 29.8 | 29760.1 |  |  |
| | | Monalisa | 100.0 | 664.9 | 39.4 | 23520.1 | $< 10^{-15}$ | 6240.1 |
|  | 2016 | Bintje | 99.8 | 473.4 | 19.1 | 42583.4 |  |  |
| | | Monalisa | 98.7 | 510.4 | 18.7 | 38453.1 | $< 10^{-15}$ | 4130.3 |
| Non irrigated |  |  |  |  |  |  |  |  |
|  | 2012 | Bintje | 97.7 | 445.8 | 11.7 | 44378.4 |  |  |
| | | Monalisa | 99.8 | 470.6 | 12.9 | 42836.1 | $< 10^{-5}$ | 1542.3 |
|  | 2013 | Bintje | 76.7 | 490.4 | 2.3 | 31404.2 |  |  |
| | | Monalisa | 60.1 | 511.2 | 12.0 | 23350.3 | $< 10^{-9}$ | 8053.9 |
|  | 2014 | Bintje | 99.4 | 689.5 | 8.9 | 20924.6 |  |  |
| | | Monalisa | 5.3 | 683.7 | 27.3 | 1145.2 | $< 10^{-15}$ | 19779.4 |

**Figure 6.**
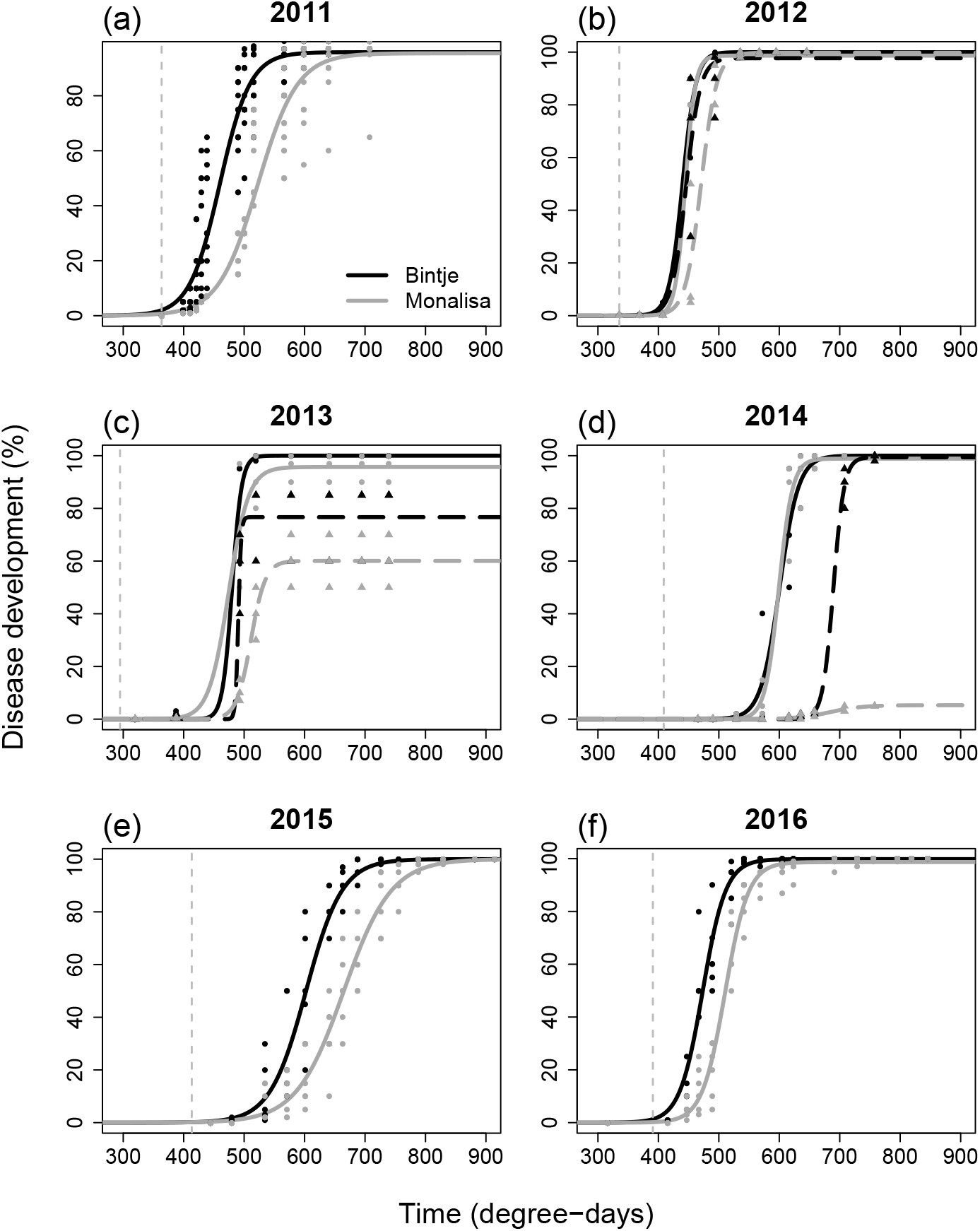
Disease progress curves. Observed data and estimated logistic curves across six years of experiments. Black lines and dots represent the Bintje variety, while grey lines and dots represent Monalisa. Solid lines indicate disease progress in irrigated plots (2011–2016), and dashed lines (2012–2014) show disease development in non-irrigated plots. For each year, the inoculation time is marked by a vertical grey dotted line.

First, *P. infestans* spread was always more severe on Bintje than on Monalisa suggesting an impact of plant architecture on disease progress. Besides the visual assessment of disease progress curves, in all cases the difference of AUDPC (benefit Δ *>* 0) in favour of stronger disease development on Bintje (Table 1). This slower development on Monalisa can be attributed to an architecture that is less favourable to the pathogen. However, the intensity of this architecture-driven disease mitigation was variable. Disease reduction was significant in all experiments without irrigation, but only in half of the irrigated experiments (Table 1).

Second, perhaps unsurprisingly, our results point out the importance of climatic conditions on architecture-driven disease mitigation. In non-irrigated plots the difference in disease development between Bintje and Monalisa was always higher than in irrigated conditions (Fig. 6 & Table 1). Moreover, as illustrated by the calculated benefit Δ, the effect of architecture was also more variable without irrigation. This is well illustrated by the contrast between 2012, where the epidemic on Monalisa was mostly delayed, and 2014 during which the disease almost did not take-off in Monalisa while it colonised all the canopy of Bintje.

### 3.4 Correlations between disease suppression and canopy architecture

Considering irrigated and non-irrigated plots together, Pearson correlation coefficients indicate that the level of disease suppression was respectively strongly and moderately associated with differences in canopy closure (*r* = 0.72) and leaf surface (*r* = 0.54) (Fig. 7). Differences in height and stem number was only weakly associated to AUDPC differences. Then the slower spread of *P. infestans* on Monalisa may be mostly explained by lower canopy closure at the time of infection, with a more limited impact of lower canopy foliage density (Table 2).

**Table 2.** Differences between canopy traits. For all traits the difference is obtain by considering Bintje minus Monalisa.

|  | Year | Height | Surface | Closure | Stems | Yield | Benefit |
| --- | --- | --- | --- | --- | --- | --- | --- |
| Irrigated |  |  |  |  |  |  |  |
|  | 2011 | -14.24 | 628.68 | -2.14 | -0.07 | -8.90 | 6167.60 |
|  | 2012 | -2.28 | -232.13 | 0.60 | 0.19 | -32.83 | 1041.50 |
|  | 2013 | 2.16 | 841.89 | 2.61 | 0.19 | -53.83 | 1246.30 |
|  | 2014 | 4.92 | -2647.60 | 0.07 | 2.05 | -33.33 | 209.10 |
|  | 2015 | -9.93 | 506.79 | -0.01 | 2.02 | -2.00 | 6240.10 |
|  | 2016 | -8.47 | 571.46 | 1.42 | 1.67 | -12.52 | 4130.30 |
| Non irrigated |  |  |  |  |  |  |  |
|  | 2012 | -1.18 | -1174.47 | -0.55 | 0.23 | -5.17 | 1542.30 |
|  | 2013 | -4.09 | 2063.58 | -0.39 | 0.23 | -26.17 | 8053.90 |
|  | 2014 | -0.93 | 1027.01 | 8.54 | 2.09 | -5.67 | 19779.40 |

**Figure 7.**
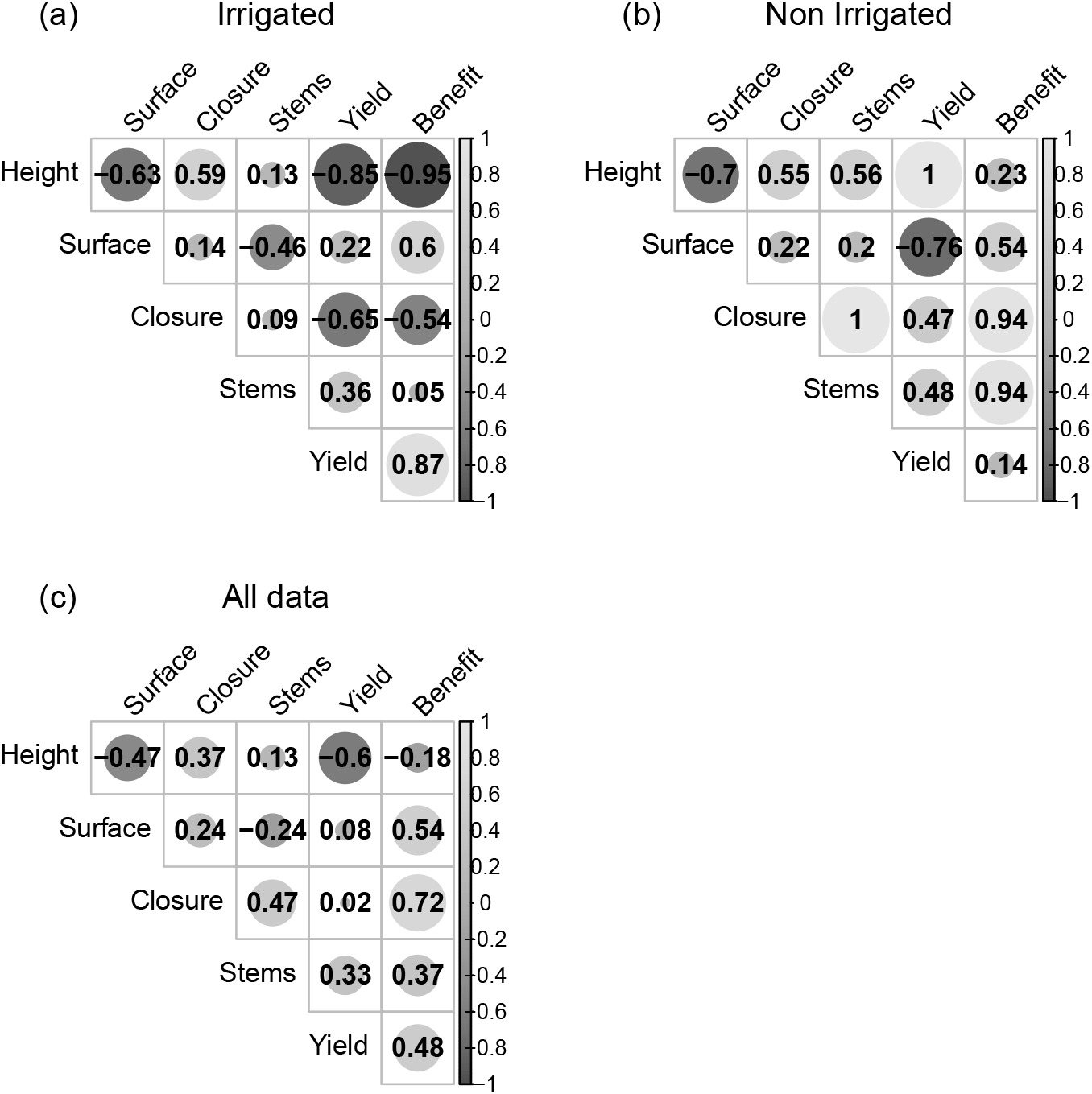
Pearson correlation coefficients among varietal differences of disease and canopy traits. Benefit ΔAUDPC between Bintje and Monalisa. Canopy closure, height and leaf surface : differences between Bintje and Mona at inoculation times.

In irrigated trials, disease suppression was most strongly correlated with increased contrasts in canopy height (*r* = *−*0.95), strongly associated with differences in leaf surface (*r* = 0.6) and moderately linked to canopy closure (*r* = *−*0.54) (Fig. 7). In this case, architecture-mediated disease mitigation in Monalisa could be explained by a higher canopy, less leaf surface and is moderately associated with a higher canopy closure when the pathogen is introduced (Table 2). This suggests that, in irrigated systems, an ‘umbrella effect’ linked to tall, porous canopies is unfavourable to late blight.

In non-irrigated trials the results suggest that disease mitigation is strongly associated with contrasts in both canopy closure (*r* = 0.94) and stem number (*r* = 0.94) with a more limited influence of leaf surface (*r* = 0.54) (Fig. 7). In less favourable environmental conditions, the less dense was the cover at pathogen arrival, the more it mitigated disease development (Table 2).

Interestingly, in irrigated trials differences in canopy height between Bintje and Monalisa was positively correlated with difference in canopy closure (*r* = 0.59) and negatively associated with contrasted leaf surface (*r* = *−*0.63). In non-irrigated plots the same trend was confirmed, and the difference in the number of stems appeared to be more strongly associated with the difference in height (*r* = 0.56) and strongly correlated with canopy closure (*r* = 1). These results confirm that Monalisa tends to grow more vertically and less horizontally than Bintje. Moreover, without additional water supply the correlations suggest that differences in canopy height might be more associated with contrasted stem numbers.

These results suggest that the disease-mitigating effect of architecture is mediated by multiple traits, with their influence varying with environmental context. Importantly, no single trait consistently predicted disease reduction across all years, reinforcing the importance of multi-year evaluations for robust inference. In addition, the analysis points out correlations between architectural traits and shows evidence that some traits, notably stem number, are influenced by water supply (Schittenhelm et al., 2006).

### 3.5 Yield analysis

Yield outcomes reflected complex interactions between cultivar, treatment, and environmental conditions (Fig. 8 & S1). Taking all the data into account together the yield was significantly affected by year, treatment and irrigation (p-values *<* 0.02) (Table S8). Unsurprisingly, the yield was higher in treated and irrigated plots.

**Figure 8.**
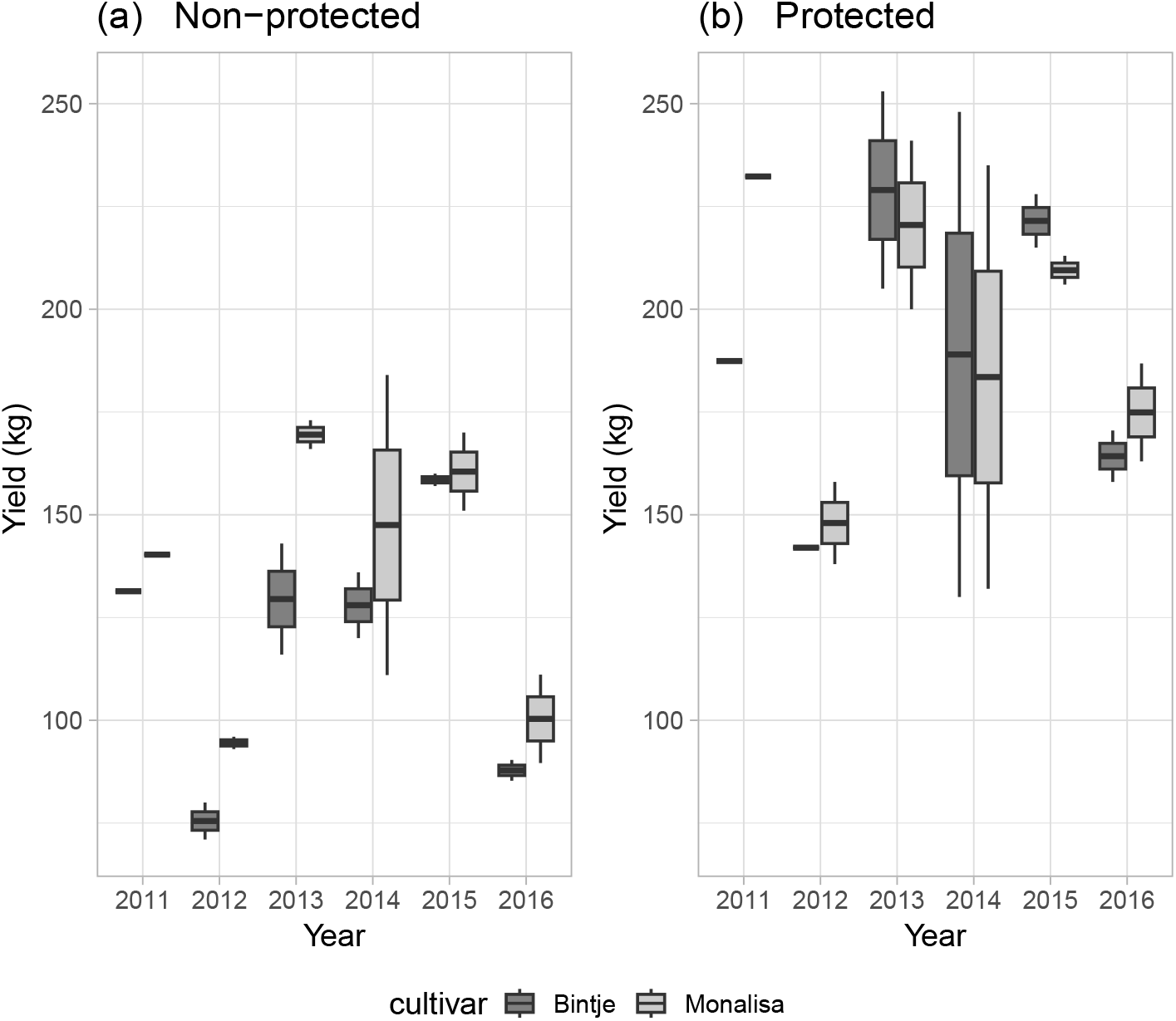
Yield data for inoculated plots. The Bintje variety is represented in dark grey, while Monalisa is in light grey.

When looking only at yield in treated trials, year and irrigation were logically the only significant factors (p-values *<* 0.02). Considering inoculated non-treated plots only, besides the year effect (p*<* 10^*−*2^) Monalisa showed marginally significant (p= 0.06) higher yields than Bintje. As illustrated by the strong correlation between AUDPC difference and yield (*r* = 0.87) under irrigated conditions, improved disease control enabled Monalisa to outperform Bintje in terms of yield (Fig. 7). This observation is quite important, as the ramose growth habit of Monalisa should have been detrimental to yield compared to the more leafy type of Bintje. This suggests that it is possible to breed for plant architectures unfavourable to late blight with no yield penalty.

In addition, the correlation analysis highlighted strong association between yield gaps and differences in architectural traits (Fig. 7). However, these correlations varied with the environmental context. For instance yield gaps were strongly associated with height contrasts but positively (*r* = 1) and negatively (*r* = *−*0.95) in respectively non-irrigated and irrigated trials. The results support the same tendency with a weaker correlation for differences in canopy closure. However, although the contrast in stem numbers was moderately associated with yield gaps, the correlation was positive in both cases (*r* = 0.36 and *r* = 0.48 respectively in irrigated and non-irrigated trials).

## 4 Discussion

This study demonstrates that potato canopy architecture can modulate the development of late blight epidemics caused by *P. infestans*, though this effect is highly variable across years and environmental conditions. By using two cultivars with comparable tissue susceptibility but contrasting architectures, we were able to focus specifically on how morphological traits influence disease dynamics under field conditions. A defining feature of this work is its long-term, replicated field trials, which replicated the variability growers encounter in practice. Conducted over six years, including irrigated trials and three additional years with non-irrigated plots, these experiments allowed us to assess how water supply, a common agricultural practice in certain growing regions, interacts with architectural traits to affect disease progression.

Our results confirm that canopy architecture in pure crop stands can partially suppress epidemic development of potato late blight. Cultivar Monalisa, characterised by an erect growth habit and reduced canopy closure, generally exhibited slower or delayed disease progression and, in some instances, lower final disease severity compared to Bintje. However, this advantage was not consistently significant: under irrigation, architecture-mediated control was effective in only half of the trials, whereas in non-irrigated conditions, it proved both more robust and more reliable (Table 1).

The multi-year dataset revealed substantial interannual variability in both trait expression (e.g. canopy height, closure, stem number) and disease outcomes, despite the stable genetic backgrounds of host cultivars (potato cultivars are vegetatively multiplicated clones). This phenotypic plasticity, likely driven by environmental interactions (Schittenhelm et al., 2006; Pangga et al., 2011), modulated the epidemiological benefits conferred by architectural traits. For example, whilst Monalisa consistently produced taller plants, the degree to which this translated into reduced disease severity depended on year-specific factors, including weather patterns or irrigation regimes (Table 2).

Critically, no single architectural variable emerged as a universal predictor of disease suppression (Fig. 7). Instead, trait-disease correlations shifted between irrigated and non-irrigated conditions: canopy height and closure were key predictors in some years, whilst leaf area and stem number played a more dominant role in others. This context dependence aligns with previous modelling and experimental studies, which highlight the multi-factorial and non-linear nature of plant disease epidemics and show that small initial changes can induce large differences in epidemic behaviour (Truscott and Gilligan, 2003; Gilligan, 2007; Pangga et al., 2011). However, these trait-disease correlations should be interpreted with caution as they rely on six (irrigated) or three (non-irrigated) year-level data points. The correlation coefficients, yet consistent in sign, are not a robust basis for ranking the relative importance of individual traits. Nevertheless the architecture-driven suppression was observed in every single trial over six years, regardless of its magnitude or statistical significance. To our knowledge, datasets that combine six years of replicated field trials with longitudinal monitoring of canopy and disease are still rare in plant disease epidemiology. Yet, even this effort falls short of what would be needed to disentangle the relative contribution of individual canopy traits with confidence.

That said, we observed a consistent impact of canopy porosity on disease reduction. In other words, short and dense canopies (like those of Bintje) are more prone to severe epidemics than taller and porous canopies, like those of Monalisa. Canopy porosity results both from plant characteristics (more erect stems and leaves, smaller leaves, fewer stems) and from crop management (planting density, irrigation, etc.) (Casadebaig et al., 2012). There is ample evidence that disease development of most aerial fungal pathogens is strongly dependent on wetness duration (Duthie, 1997; Richard et al., 2013). As more porous canopies tend to dry faster after rainfall or irrigation, their more limited conduciveness to severe epidemic development probably results from the impact of canopy architecture on crop microclimate (Tivoli et al., 2013). Part of it might also result from differences in inoculum deposition to susceptible organs, but this would require additional experiments to ascertain. An important observation was that disease mitigation did not always translate into yield benefits (Fig. 8 & Table S8). Whilst Monalisa occasionally outperformed Bintje in irrigated plots during years with strong disease suppression, yield advantages remained modest and inconsistent. Under fungicide protection, yield differences between cultivars were negligible, confirming that the observed effects were disease-mediated rather than inherent to productivity potential. This decoupling between disease control and yield benefit echoes other studies that manipulated potato canopy or crop configuration against late blight (Rakotonindraina et al., 2012). For instance, Hospers-Brands et al. (2008) found that varying plant population and spacing in organic potato crops had no measurable effect on late blight infection, yet still resulted in significant, disease-independent effects on tuber size grading. Conversely, Bouws and Finckh (2008) showed that strip-cropping potato with cereals reduced incoming inoculum without changing the underlying disease-yield-loss relationship. Together, these examples illustrate that architectural or agronomic levers can affect yield and disease along largely independent pathways, so that a benefit on one axis need not translate into a benefit on the other, in either direction.

These findings show that while canopy characteristics can and should be considered in integrated crop management strategies, they are not sufficient alone to guarantee a high level of disease control or a yield benefit. Therefore, architectural traits can complement other control measures, such as genetic resistance or fungicide applications, and need to be integrated into holistic, context-specific disease management programmes. It would be worth investigating how the partial control provided by architecture could be complemented by other control measures, such as genetic resistance or cultivar mixture (Andrivon et al., 2003; Skelsey et al., 2005; Bouws and Finckh, 2008; Hospers-Brands et al., 2008; Homulle et al., 2025). Such more integrated approaches may lead to redesigning disease-resilient crop ideotypes that effectively combine architectural, genetic, and ecological strategies for the sustainable suppression of major diseases such as potato late blight (Andrivon et al., 2013; Debaeke et al., 2021).

Our multi-year data exhibited the fluctuating nature of architectural effects and their dependence on environmental conditions. Although the mechanisms linking architecture to disease progression are well documented (Ando et al., 2007; Tivoli et al., 2013), predicting outcomes under real-world uncertainty remains a significant challenge. Implementing further multi-year and multi-environment experiments and using advanced analytical tools, including functional data analysis (Boschi et al., 2021; Matsui and Mochida, 2024) and dynamic modelling (Casadebaig et al., 2012; Mammeri et al., 2014), would help disentangle the overlapping influences of environmental, architectural and inoculum factors, thereby improving the predictability of disease outcomes under variable conditions (Pangga et al., 2011). Moreover, exploring potential feedback loops between disease progression and canopy architecture, particularly how severe defoliation might reshape canopy structure to create less favourable conditions for pathogens, represents a critical but understudied area that could refine our understanding of pathogen-host dynamics. Developing and fitting dynamic models to experimental data may help to identify if such feedback processes occur in such plant-pathogen systems (Bailey and Gilligan, 2004; Motisi et al., 2019).

## 5 Acknowledgments

This study was funded by the French National Research Agency (ANR) and the European Union’s Horizon 2020 research and innovation programs, respectively through the ARCHIDEMIO (grant ANR-08-STRA-04) and Organic-Plus (grant agreement 774340) projects. The authors warmly thank UE La Motte (INRAE, Le Rheu) for hosting the trials and conducting all cultural operations and Hervé Douchy for his valuable help.

