## Supplementary materials for "Plant architecture as a tool to mitigate late blight: significant but variable contributions of erect, aerated potato canopies over six years of field trials"

### S1 Key dates for crop management and summary of measured traits during canopy phenotyping

Table S1: Timing of main action on potato field trials.

| Year | Planting | Inoculation | Harvesting | Irrigation |
|---|---|---|---|---|
| 2011 | $12^{th}$ April | $16^{th}$ June | $19^{th}$ September | Yes |
| 2012 | $12^{th}$ April | $18^{th}$ June | $04^{th}$ September | Yes & No |
| 2013 | $22^{nd}$ April | $21^{st}$ June | $16^{th}$ September | Yes & No |
| 2014 | $11^{th}$ April | $20^{th}$ June | $08^{th}$ September | Yes & No |
| 2015 | $14^{th}$ April | $19^{th}$ June | $07^{th}$ September | Yes |
| 2016 | $18^{th}$ April | $22^{nd}$ June | $12^{th}$ September | Yes |

Table S2: Summary of measured traits.

| Trait | Treatment | Monitoring | Unit |
|---|---|---|---|
| Disease incidence | Inoculated | Longitudinal | % |
| Canopy Coverage | Inoculated | Longitudinal | % |
| Canopy Height | Inoculated | Longitudinal | cm |
| Leaf surface per plant | Protected | Longitudinal | $cm^2$ |
| Number of stems per plant | Protected | Once after inoculation | stems |
| Yield | Inoculated & Protected | Once after harvest | kg |

### S2 Canopy height

Table S3: Estimated parameters of the model for the height of inoculated canopies

| | Year | Cultivar | $\kappa$ | $\beta$ | $\gamma$ | $\delta$ | $\nu$ |
|---|---|---|---|---|---|---|---|
| Irrigated | | | | | | | |
| | 2011 | Bintje | 37.0 | 16.4 | 75.6 | 574.6 | 34.0 |
| | | Monalisa | 53.4 | 98.6 | 67.5 | 482.7 | 34.0 |
| | 2012 | Bintje | 96.2 | 183.2 | 73.0 | 674.5 | 169.4 |
| | | Monalisa | 121.3 | 150.6 | 75.6 | 503.8 | 188.5 |
| | 2013 | Bintje | 105.0 | 183.5 | 61.0 | 754.4 | 206.8 |
| | | Monalisa | 92.5 | 144.5 | 39.4 | 754.4 | 248.0 |
| | 2014 | Bintje | 71.7 | 166.9 | 49.3 | 613.2 | 24.2 |
| | | Monalisa | 66.6 | 161.3 | 45.1 | 615.5 | 27.3 |
| | 2015 | Bintje | 65.7 | 149.9 | 54.1 | 697.5 | 34.0 |
| | | Monalisa | 77.3 | 183.6 | 65.1 | 697.5 | 34.0 |
| | 2016 | Bintje | 57.1 | 155.7 | 50.4 | 545.5 | 10.0 |
| | | Monalisa | 115.5 | 175.3 | 79.7 | 551.3 | 311.8 |
| Non irrigated | | | | | | | |
| | 2012 | Bintje | 76.4 | 172.2 | 81.9 | 909.4 | 165.8 |
| | | Monalisa | 123.6 | 150.5 | 102.8 | 503.9 | 228.2 |
| | 2013 | Bintje | 148.0 | 251.3 | 90.1 | 518.8 | 136.2 |
| | | Monalisa | 86.8 | 150.1 | 90.1 | 525.2 | 47.2 |
| | 2014 | Bintje | 50.7 | 136.4 | 57.6 | 999.7 | 352.1 |
| | | Monalisa | 44.9 | 130.4 | 51.2 | 903.6 | 148.0 |

### S3 Canopy closure

Table S4: Estimated parameters for inoculated canopy coverage. The carrying capacity parameter $\kappa$ is fixed to 100.

| | Year | Cultivar | $\beta$ | $\gamma$ | $\delta$ | $\nu$ |
|---|---|---|---|---|---|---|
| Irrigated | | | | | | |
| | 2011 | Bintje | 103.2 | 85.6 | 456.1 | 31.4 |
| | | Monalisa | 137.6 | 77.2 | 518.8 | 41.4 |
| | 2012 | Bintje | 49.9 | 18.7 | 448.3 | 20.6 |
| | | Monalisa | 49.7 | 21.3 | 452.2 | 25.5 |
| | 2013 | Bintje | 75.3 | 35.3 | 497.3 | 38.5 |
| | | Monalisa | 59.7 | 28.6 | 511.3 | 64.1 |
| | 2014 | Bintje | 117.8 | 34.5 | 606.5 | 13.6 |
| | | Monalisa | 123.8 | 41.0 | 606.5 | 14.4 |
| | 2015 | Bintje | 105.5 | 33.0 | 640.7 | 23.9 |
| | | Monalisa | 124.7 | 30.6 | 702.8 | 27.0 |
| | 2016 | Bintje | 134.6 | 41.7 | 504.4 | 19.6 |
| | | Monalisa | 151.7 | 47.1 | 570.9 | 41.7 |
| Non irrigated | | | | | | |
| | 2012 | Bintje | 62.0 | 50.4 | 456.2 | 22.3 |
| | | Monalisa | 53.5 | 47.5 | 479.0 | 19.7 |
| | 2013 | Bintje | 91.8 | 37.8 | 547.0 | 67.8 |
| | | Monalisa | 52.2 | 39.8 | 537.8 | 64.2 |
| | 2014 | Bintje | 78.6 | 172.2 | 688.3 | 13.5 |
| | | Monalisa | 142.9 | 92.4 | 615.4 | 124.7 |

### S4 Leaf surface per plant

Table S5: Estimated parameters of the model for the leaf surface on protected canopies

| | Year | Cultivar | $\kappa$ | $\beta$ | $\gamma$ | $\delta$ | $\nu$ |
|---|---|---|---|---|---|---|---|
| Irrigated | | | | | | | |
| | 2011 | Bintje | 9000.0 | 175.8 | 179.9 | 1000.0 | 301.0 |
| | | Monalisa | 8500.0 | 175.8 | 218.2 | 1000.0 | 301.0 |
| | 2012 | Bintje | 12748.9 | 156.1 | 77.3 | 842.6 | 330.2 |
| | | Monalisa | 11664.2 | 135.7 | 68.3 | 842.6 | 263.8 |
| | 2013 | Bintje | 14140.3 | 146.7 | 62.5 | 842.6 | 324.9 |
| | | Monalisa | 12035.9 | 138.2 | 41.8 | 842.6 | 324.9 |
| | 2014 | Bintje | 8971.8 | 180.0 | 100.1 | 1000.0 | 301.0 |
| | | Monalisa | 16026.5 | 211.4 | 69.3 | 527.6 | 180.0 |
| | 2015 | Bintje | 14258.1 | 160.0 | 53.4 | 791.7 | 372.1 |
| | | Monalisa | 34813.8 | 77.5 | 118.9 | 10.0 | 602.8 |
| | 2016 | Bintje | 20000.0 | 305.1 | 141.3 | 617.0 | 100.0 |
| | | Monalisa | 14931.3 | 237.7 | 69.6 | 638.3 | 176.8 |
| Non irrigated | | | | | | | |
| | 2012 | Bintje | 39892.1 | 391.6 | 120.8 | 490.7 | 83.6 |
| | | Monalisa | 214481.2 | 411.3 | 105.9 | 432.9 | 101.1 |
| | 2013 | Bintje | 176281.0 | 169.9 | 136.3 | 200.8 | 179.4 |
| | | Monalisa | 13773.9 | 171.6 | 40.3 | 497.6 | 31.3 |
| | 2014 | Bintje | 14771.9 | 144.1 | 48.2 | 338.1 | 229.9 |
| | | Monalisa | 10445.7 | 156.9 | 76.0 | 432.0 | 192.4 |

### S5 Statistical analysis of the number of stems in field trials.

Table S6: ANOVA table for assessing the effects of cultivar, year and irrigation on the number of stems.

| Effect | Sum of squares | Df | F value | Pr(>F) |
|---|---|---|---|---|
| Cultivar | 228.63 | 1 | 81.09 | $\mathbf{< 10^{-15}}$ |
| Year | 245.39 | 5 | 17.41 | $\mathbf{< 10^{-15}}$ |
| Irrigation | 1.33 | 1 | 0.47 | 0.49 |
| Cultivar × Year | 74.31 | 5 | 5.27 | $\mathbf{< 10^{-3}}$ |
| Cultivar × Irrigation | 0.02 | 1 | 0.01 | 0.93 |

Table S7: Estimated marginal means for the number of stems

| Cultivar | Least-squares means | SE | df | lower.CL | upper.CL | Group |
|---|---|---|---|---|---|---|
| *2011 - Irrigated* | | | | | | |
| Bintje | 3.7333 | 0.4335 | 448 | 2.8813 | 4.5854 | 1 |
| Monalisa | 3.8000 | 0.4335 | 448 | 2.9480 | 4.6520 | 1 |
| *2012 - Irrigated* | | | | | | |
| Monalisa | 6.8199 | 0.3606 | 448 | 6.1112 | 7.5287 | 1 |
| Bintje | 7.0134 | 0.3606 | 448 | 6.3046 | 7.7221 | 1 |
| *2013 - Irrigated* | | | | | | |
| Monalisa | 5.3438 | 0.3832 | 448 | 4.5907 | 6.0968 | 1 |
| Bintje | 5.5312 | 0.3832 | 448 | 4.7782 | 6.2843 | 1 |
| *2014 - Irrigated* | | | | | | |
| Monalisa | 3.8816 | 0.3057 | 448 | 3.2809 | 4.4824 | 1 |
| Bintje | 5.9290 | 0.3057 | 448 | 5.3282 | 6.5297 | 2 |
| *2015 - Irrigated* | | | | | | |
| Monalisa | 3.6029 | 0.2036 | 448 | 3.2028 | 4.0031 | 1 |
| Bintje | 5.6176 | 0.2036 | 448 | 5.2175 | 6.0178 | 2 |
| *2016 - Irrigated* | | | | | | |
| Monalisa | 3.8077 | 0.2328 | 448 | 3.3501 | 4.2653 | 1 |
| Bintje | 5.4808 | 0.2328 | 448 | 5.0232 | 5.9384 | 2 |
| *2011 - Non irrigated* | | | | | | |
| Bintje | 3.9208 | 0.5527 | 448 | 2.8347 | 5.0070 | 1 |
| Monalisa | 3.9458 | 0.5527 | 448 | 2.8597 | 5.0320 | 1 |
| *2012 - Non irrigated* | | | | | | |
| Monalisa | 6.9658 | 0.3606 | 448 | 6.2570 | 7.6745 | 1 |
| Bintje | 7.2009 | 0.3606 | 448 | 6.4921 | 7.9096 | 1 |
| *2013 - Non irrigated* | | | | | | |
| Monalisa | 5.4896 | 0.3832 | 448 | 4.7365 | 6.2427 | 1 |
| Bintje | 5.7187 | 0.3832 | 448 | 4.9657 | 6.4718 | 1 |
| *2014 - Non irrigated* | | | | | | |
| Monalisa | 4.0275 | 0.3057 | 448 | 3.4267 | 4.6282 | 1 |
| Bintje | 6.1165 | 0.3057 | 448 | 5.5157 | 6.7172 | 2 |
| *2015 - Non irrigated* | | | | | | |
| Monalisa | 3.7488 | 0.3987 | 448 | 2.9653 | 4.5323 | 1 |
| Bintje | 5.8051 | 0.3987 | 448 | 5.0217 | 6.5886 | 2 |
| *2016 - Non irrigated* | | | | | | |
| Monalisa | 3.9535 | 0.4144 | 448 | 3.1392 | 4.7679 | 1 |
| Bintje | 5.6683 | 0.4144 | 448 | 4.8539 | 6.4826 | 2 |

Confidence level used: 0.95

significance level used: alpha = 0.05

### S6 Statistical analysis of yield data

Table S8: ANOVA table for assessing the effects of year, cultivar, treatment and irrigation on yield data.

| Effect | Sum of squares | Df | F value | Pr(>F) |
|---|---|---|---|---|
| All data | | | | |
| Year | 34694.24 | 5 | 10.37 | $\mathbf{< 10^{-3}}$ |
| Cultivar | 1111.53 | 1 | 1.66 | 0.21 |
| Treatment | 45116.82 | 1 | 67.46 | $\mathbf{< 10^{-6}}$ |
| Irrigation | 4620.37 | 1 | 6.91 | **0.02** |
| Year × Cultivar | 895.10 | 5 | 0.27 | 0.92 |
| Year × Treatment | 1545.42 | 5 | 0.46 | 0.79 |
| Cultivar × Treatment | 647.19 | 1 | 0.97 | 0.34 |
| Cultivar × Irrigation | 70.04 | 1 | 0.10 | 0.75 |
| Treatment × Irrigation | 2542.04 | 1 | 3.80 | **0.07** |
| Year × Cultivar × Treatment | 1945.16 | 5 | 0.58 | 0.71 |
| Cultivar × Treatment × Irrigation | 651.04 | 1 | 0.97 | 0.34 |
| Inoculated | | | | |
| Cultivar | 1727.52 | 1 | 4.72 | **0.06** |
| Year | 18133.28 | 5 | 9.91 | $\mathbf{< 10^{-2}}$ |
| Irrigation | 154.08 | 1 | 0.42 | 0.53 |
| Cultivar × Year | 1328.38 | 5 | 0.73 | 0.62 |
| Cultivar × Irrigation | 574.08 | 1 | 1.57 | 0.24 |
| Protected | | | | |
| Cultivar | 31.20 | 1 | 0.03 | 0.86 |
| Year | 18106.37 | 5 | 3.73 | **0.048** |
| Irrigation | 7008.33 | 1 | 7.21 | **0.027** |
| Cultivar × Year | 1511.87 | 5 | 0.31 | 0.89 |
| Cultivar × Irrigation | 147.00 | 1 | 0.15 | 0.70 |

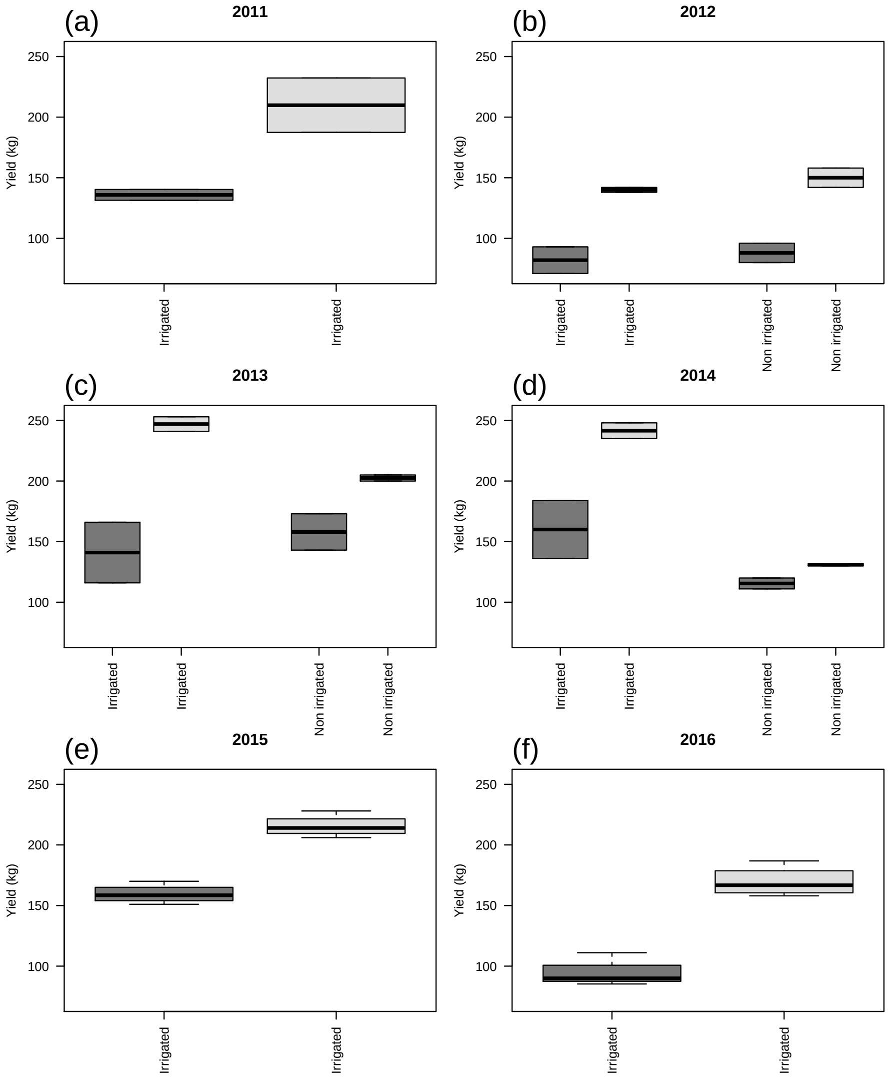


Figure S1: Distribution of yield data across experimental conditions.
